# Target class profiling of small molecule methyltransferases

**DOI:** 10.1101/2022.03.24.485659

**Authors:** Quinlin M Hanson, Min Shen, Hui Guo, Ig-Jun Cho, Matthew D Hall

## Abstract

Target class profiling (TCP) is a chemical biology approach to investigating understudied biological target classes. TCP is achieved by developing a generalizable assay platform and screening curated compound libraries to interrogate the chemical biological space of members of an enzyme family with clinical and biological importance. In this work we took a TCP approach to investigate inhibitory activity across a set of small molecule methyltransferases, a subclass of methyltransferase enzymes, with the goal of creating a launchpad to explore this largely understudied target class. Using the representative enzymes nicotinamide N-methyltransferase (NNMT), phenylethanolamine N-methyltransferase (PNMT), histamine N-methyltransferase (HNMT), glycine N-methyltransferase (GNMT), catechol O-methyltransferase (COMT), and guanidinoacetate N-methyltransferase (GAMT), we optimized high-throughput screening (HTS)-amenable assays to screen 27,574 unique small molecules against all targets. From this dataset we identified a novel inhibitor which selectively inhibits the small molecule methyltransferase HNMT, and demonstrated how this platform approach can be leveraged for a targeted drug discovery campaign using the example of HNMT.

## Introduction

Target class profiling (TCP) is a chemical biology approach for probe development, which is complimentary to traditional disease-focused drug discovery ^1^. There are several key differences in the goals and outcomes of target class profiling versus disease-focused drug discovery. Target class profiling takes a broad approach by simultaneously studying multiple members of an enzyme family with clinical and biological relevance, whereas disease-focused drug discovery is a high-risk, high-reward endeavor focused on a single target ^2, 3^. The desired outcome for target class profiling is generation of specialized knowledge for a target class through structure-activity relationship (SAR) profiling using defined compound libraries, while disease-focused drug discovery aims for a clinically viable and marketable compound. Effectively, target class profiling emphasizes platform creation for a family of proteins, agnostic of a specific target, over product development.

The target agnostic nature of target class profiling has two key benefits. First, this philosophy makes target class profiling amenable to including understudied targets within the target class study, which aligns with efforts such as Target 2035 to create probes (tool molecules) for all human proteins ^4^. Second, the SAR profiles generated can be leveraged as a launchpad to jumpstart multiple probe development or target-focused drug discovery campaigns within the target class. A target class is a family of enzymes which are structurally and functionally related^1^. The best-studied target classes are kinases^5–7^ and GPCRs^8, 9^. Methyltransferases (MTases) in general are considered an ‘emerging’ target class with small molecule methyltransferases (SMMTases) constituting an understudied subset of this target class^2, 10–14^. In this work we apply target class profiling to SMMTase with the goal of creating a launchpad for methyltransferase probe development.

MTases are a family of enzymes which catalyze the transfer of a methyl group from a cofactor (typically S-adenosylmethionine, SAM) to a protein, nucleic acid, or small molecule substrate, with the concomitant production of S-adenosyl homocysteine (SAH) ^13, 15^. MTases are functionally classified by the atom of the substrate to which the methyl group is transferred. The most common types of MTases are *O*- and *N*-MTases, which affix methyl groups to oxygen and nitrogen atoms, respectively, with *S*-, *C*-, and metal MTases also existing. ^12, 16^. SMMTases primarily include O- and N-MTases, but some S- and C-MTases have been described. Across the human methyltransferome, SMMTases are found scattered about the various MTase clades, suggesting substrate type-preference is disjointed from primary amino acid sequence (Figure 1A) ^17, 18^. The core catalytic domain tertiary structure of SMMTases is conserved, primarily falling into the Rossman-like fold of Class I MTases^12, 16, 19^. The major structural differences of SMMTases occur in the substrate binding pocket where solvent accessibility and binding pocket volume are varied (Figure 1C) or in structural additions inserted into loops, which contribute to active site accessibility. The SAM-binding pocket for MTases is highly conserved as is the catalytic mechanism of methyl transfer^15^.

**Figure 1.**
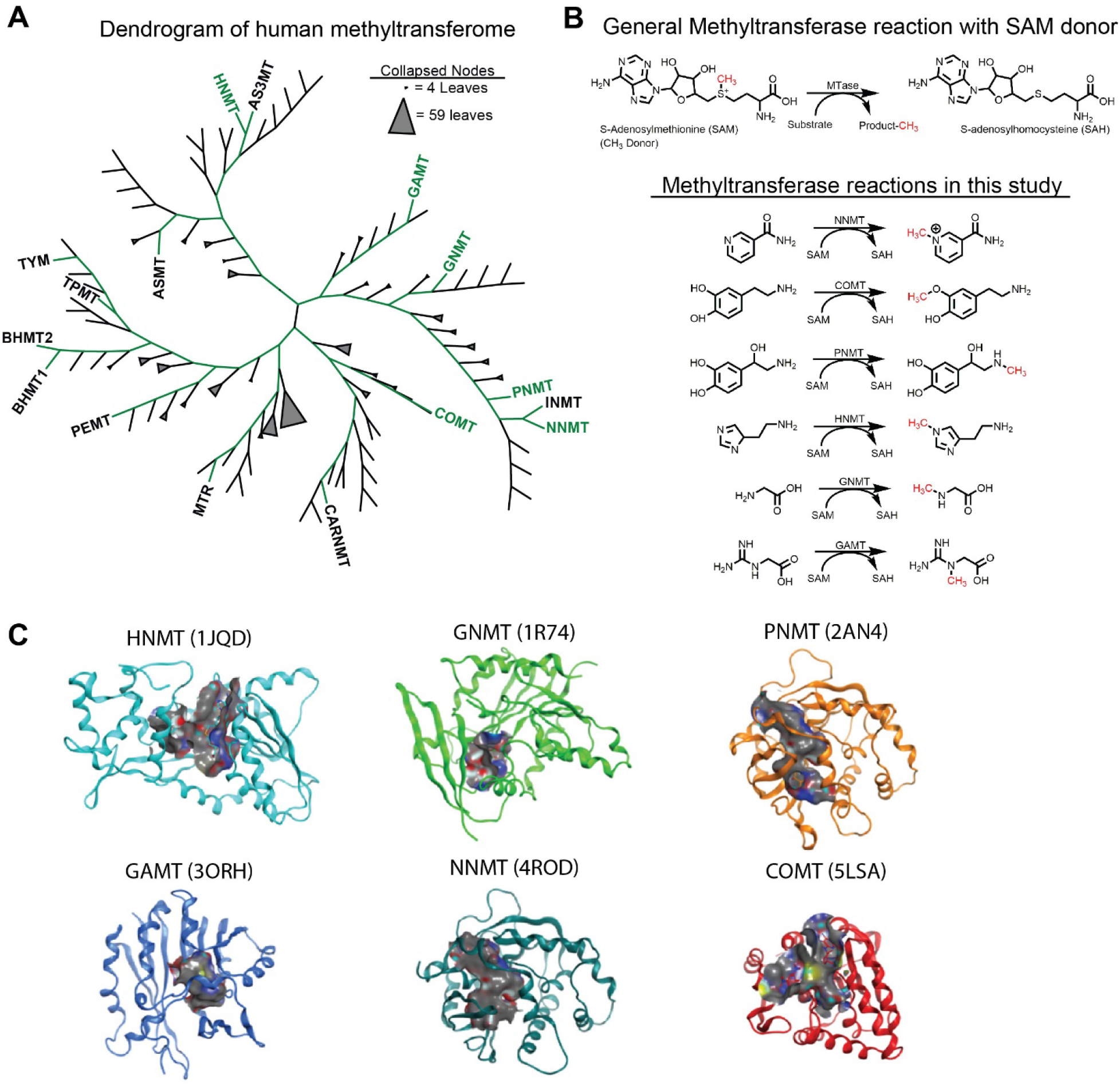
Small molecule methyltransferases comprise a diverse subset of methyltransferases. A) Dendrogram of human methyltransferases. Methyltransferase sequences were retrieved from UniProt and aligned using ClustalW. Dendrogram image was produces using ITOL. Larger nodes were collapsed into triangles. Green paths lead to small molecule methyltransferases. MTase names highlighted in green indicate those used in this study. B) General reaction diagram for methyltransferases which use SAM as a methyl donor. Also shown are biochemical reactions for the SMMTases used in this study: NNMT, COMT, PNMT, HNMT, GNMT, and GAMT. C) Structures of SMMTases reveal a large variation in solvent-accessible space (grey surfaces) among the representative enzymes used in this study. Alpha carbon RMSD for all structures is 1.59 Å. PDB IDs: HNMT (1JQD), GNMT (1R74), PNMT (2AN4), GAMT (3ORH), NNMT (4ROD), COMT (5LSA).

MTases have long been an enticing target class because of their association with diseases ranging from cancer to neurological disorders to metabolic syndromes ^2, 10, 11, 17, 19, 20^. Most MTases act as epigenetic regulators, either by modifying protein and nucleic acid substrates to directly modulate gene accessibility, or alternatively via consumption of SAM thus depleting the cellular methyl pool available ^15, 17, 19, 21–23^. SMMTases act on small molecule substrates (also referred to as metabolic substrates) to influence signaling pathways or metabolic flux. The best-known SMMTase is perhaps catecholamine O-methyltransferase (COMT), which methylates catechol-containing small molecules to terminate adrenaline signaling pathways and metabolize drugs ^20, 24–27^. Inhibition of COMT activity by the FDA-approved drugs tolcapone and entacapone is a key component of combinatorial drug therapies for Parkinson’s disease^20, 25^. Other notable functions of SMMTases include the termination of adrenaline signaling pathways by phenolethanolamine N-methyltransferase (PNMT) ^28, 29^, detoxifying xenobiotics (drug metabolism) by thiopurine S-methyltransferase (TPMT) ^30, 31^, and managing the available methyl pool for all MTase activity by glycine N-methyltransferase (GNMT)^32, 33^. Despite their central roles in biological processes, SMMTases constitute a target class of viable and largely understudied therapeutic targets.

In this work we created a target class profiling platform for SMMTases. Enzymatic assays were developed for PNMT, nicotinamide N-methyltransferase (NNMT), COMT, histamine N-methyltransferase (HNMT), GNMT, and guanidinoacetate N-methyltransferase (GAMT), adapted to 1536-well plate assays for automated screening, and all six assays were screened against 25,574 compounds. This set of compounds was refined to 1,434 active compounds (against at least one SMMTase), which had their activity confirmed across the target class and a counter-assay. From these 1,434 compounds we were able to identify a pan-active chemotype (putative tool molecule) and an HNMT-selective chemotype which typify the insight derived from target class profiling.

## Results and Discussion

### HTS Assay Development

Creating an assay platform for target class profiling required selection of representative members of the target class. We were able to source and confirm enzymatic activity for the SMMTases NNMT, PNMT, COMT, HNMT, GNMT, and GAMT, and selected them as representative SMMTases. Each of these SMMTases use SAM as a methyl donor cofactor but act on a variety of biological substrate (Figure 1B). The solvent accessible space in the active site widely varies across these six enzymes (Figure 1C).

To develop a generalizable HTS assay platform we used the MTase-Glo kit (Promega) to measure the production of SAH, a common product of all MTases being studied. The MTase-Glo system is a proprietary coupled-enzyme reagent that converts SAH to ATP through a series of enzymatic reactions including SAH hydrolase, polyphosphate-AMP phosphotransferase, and adenosine kinase^34^. ATP generation is monitored using a luciferase/luciferin bioluminescent reaction, ultimately generating a luminescent signal proportional to the SAH concentration and therefore MTase enzymatic activity. For each enzyme we determined the optimal biochemical assay conditions according to standard HTS practices^35^. K_M_ values for each substrate and the SAM cofactor were determined using the method of initial rates (Figure 2A, B, Table 2). Each of the determined values were comparable to published K_M_ values^28, 36–41^. We determined the 20% product formation time for each assay using substrate and cofactor at K_M_ concentrations to select an assay time window (PNMT example in Figure 2C). To ensure the assays were sensitive to inhibition, we tested each SMMTase against the non-specific MTase inhibitor sinefungin in 11-point dose response (Figure 2D). All MTases except GNMT were inhibitable at concentrations tested (up to 50 μM). A MTase-Glo counter-assay was developed to identify false-positive hits inhibiting the coupled reporter assay. The counter-assay was developed by directly adding 1 μM SAH (correlating to 10-20% product conversion) to the MTase-Glo reaction in place of an enzyme, and the counter-assay was not inhibited by sinefungin.

**Figure 2.**
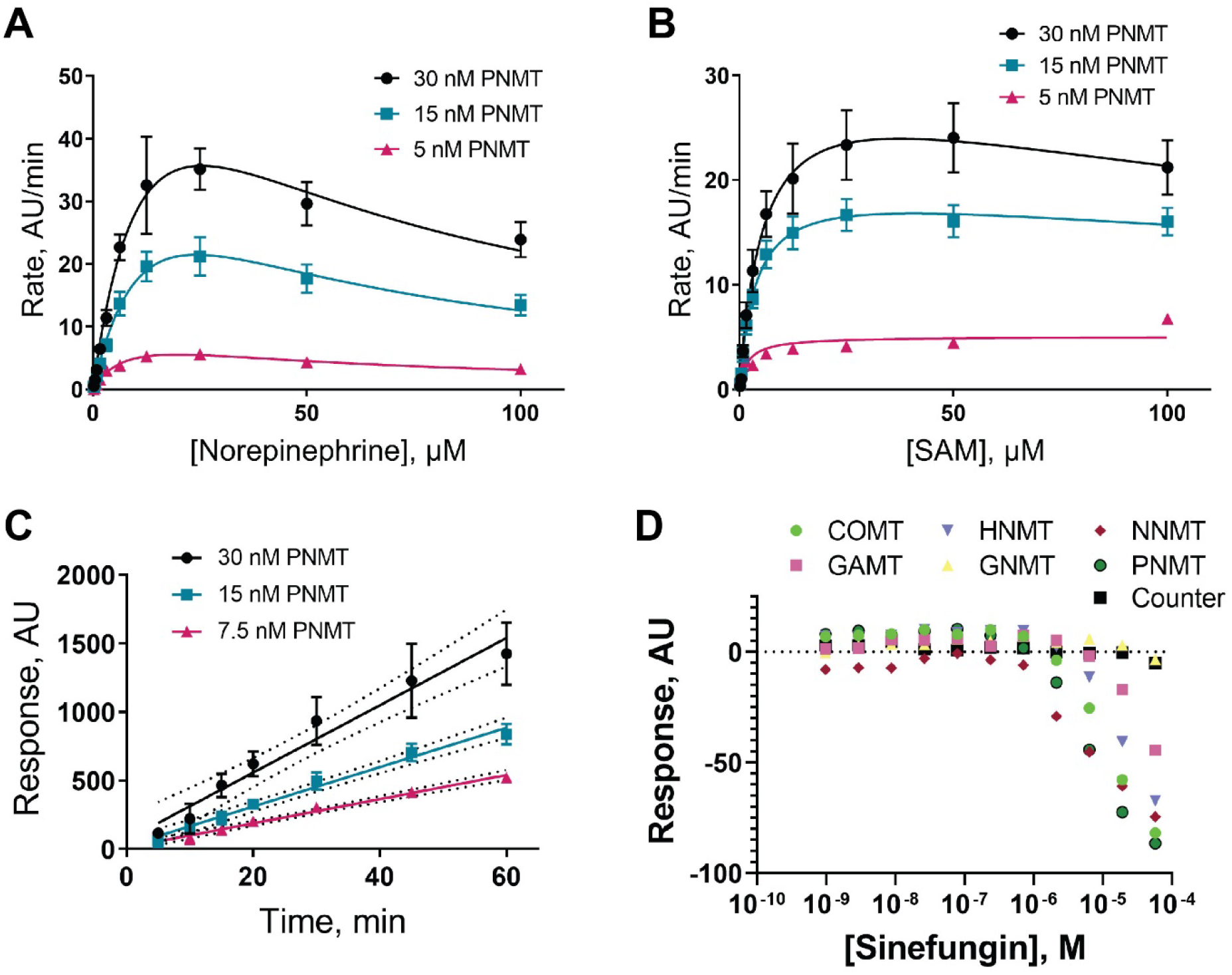
Biochemical assay development for PNMT shown as a representative example. A) K_M_ for PNMT substrate norepinephrine was determined using the method of initial rates to determine Michaelis constants. Data were plotted and fitted to a substrate inhibition model using GraphPad prism. SAM was held constant at 100 uM. B) K_M_ for SAM was determined as described in A. Norepinephrine was held constant at 100 μM. C) Reaction time course at 30 nM, 15 nM, and 7.5 nM PNMT were performed with SAM and norepinephrine at K_M_ concentrations to determine when 10-20% conversion was achieved. At 30 nM PNMT the reaction begins to plateau at 50 min and 20% conversion was estimated at 20 min. D) All SMMTases used in this project were tested against sinefungin, a non-specific positive control MTase inhibitor.

**Table 1:**
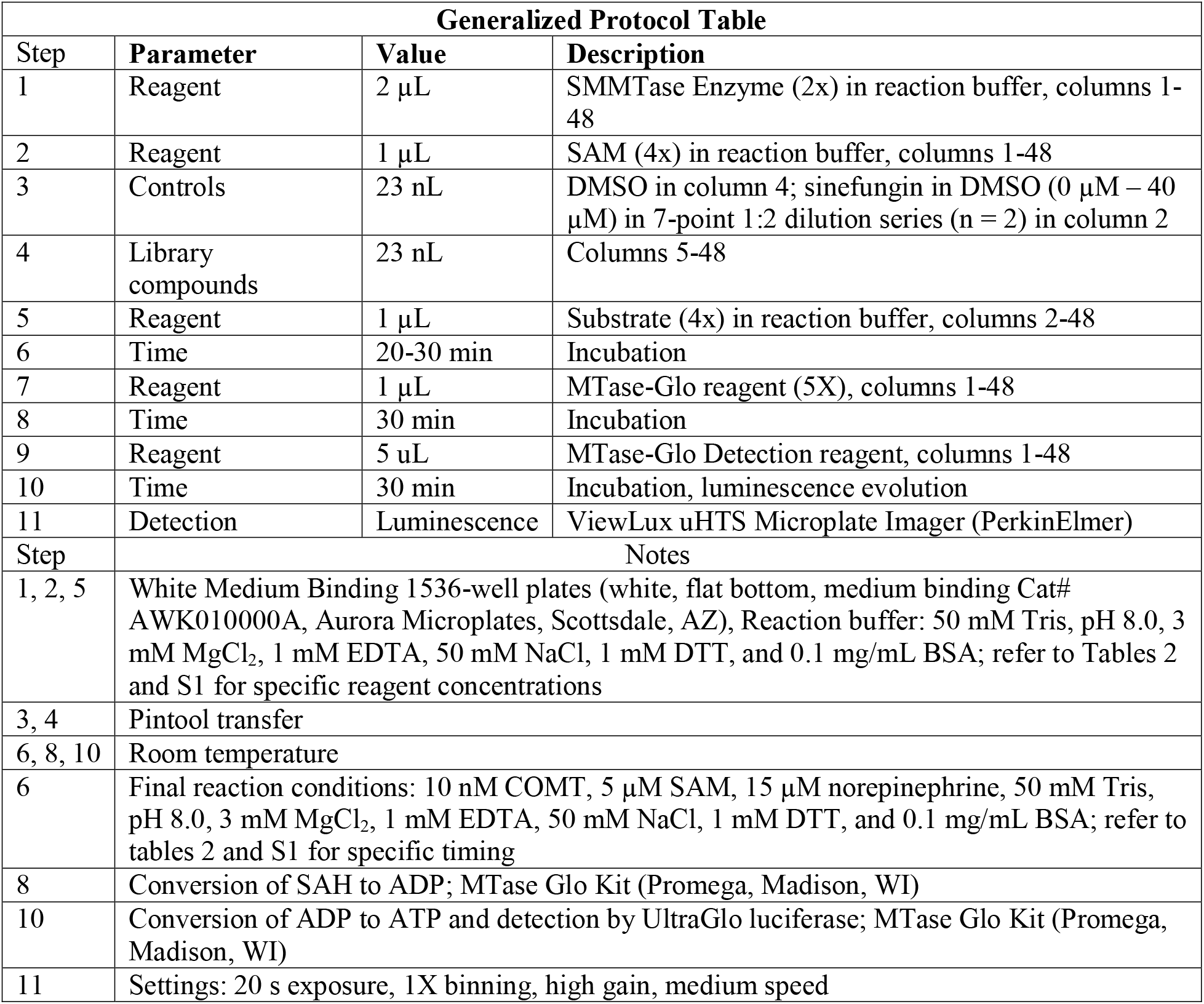
Generalized protocol table summarizing HTS assays.

**Table 2.** Michaelis-Menten constants were determined for the substrates and SAM cofactor of HNMT, GNMT, PNMT, COMT, NNMT, and GAMT using the method of initial rates.

| **SMMTase** | **Substrate** | **K<sub>M</sub>, substrate, μM** | **K<sub>M</sub>, SAM, μM** |
| --- | --- | --- | --- |
| HNMT | Histamine | 5.3±2.1 | 9.3±2.1 |
| GNMT | Glycine | >500 | 12.1±3.6 |
| PNMT | Norephinephrine | 16.2±4.3 | 3.6±0.6 |
| COMT | Norephinephrine | 13.7±2.2 | 14.5±1.8 |
| NNMT | Nicotinamide | 3.8±0.5 | 11.9±5.7 |
| GAMT | Guanidinoacetate | 2.5±0.6 | 5.6±0.8 |

### Compound Set Selection

Selecting and characterizing the appropriate compound libraries is key to developing a target class profile^1, 5^. The compound libraries tested need to balance chemical diversity and chemical relatedness. To this end we selected two diversity libraries (used to identify starting points for medicinal chemistry programs) in our collection based on the chemical space sampled and physical properties of the compound sets (Figure 3). We also selected two annotated libraries containing compounds with known pharmacologic mechanisms of action to approach the concepts of “relatedness and diversity” from a biological perspective. The chemical diversity libraries selected were the internal Genesis MiniMe collection (9,856 compounds) and a commercial MTase Library (11,000 compounds, ChemDiv, San Diego, CA). The annotated libraries we used were developed in-house: NPACT (5,099 compounds), a collection of pharmacologically active compounds and natural products, and LOPAC-Epi (1,619 compounds), a combination of LOPAC 1280 compounds that are most commonly used to validate new drug discovery assays and characterize orphan receptors, with the addition of 339 epigenetic inhibitors. The resulting collection consisted of 27,574 compounds selected for efficient and robust coverage of chemical and biological properties. The compounds selected cover a wide range of chemical space (Figure 3A) despite the majority of compounds aligning with Lipinki’s “rule of 5s”^42^ (Figure 3B-D).

**Figure 3.**
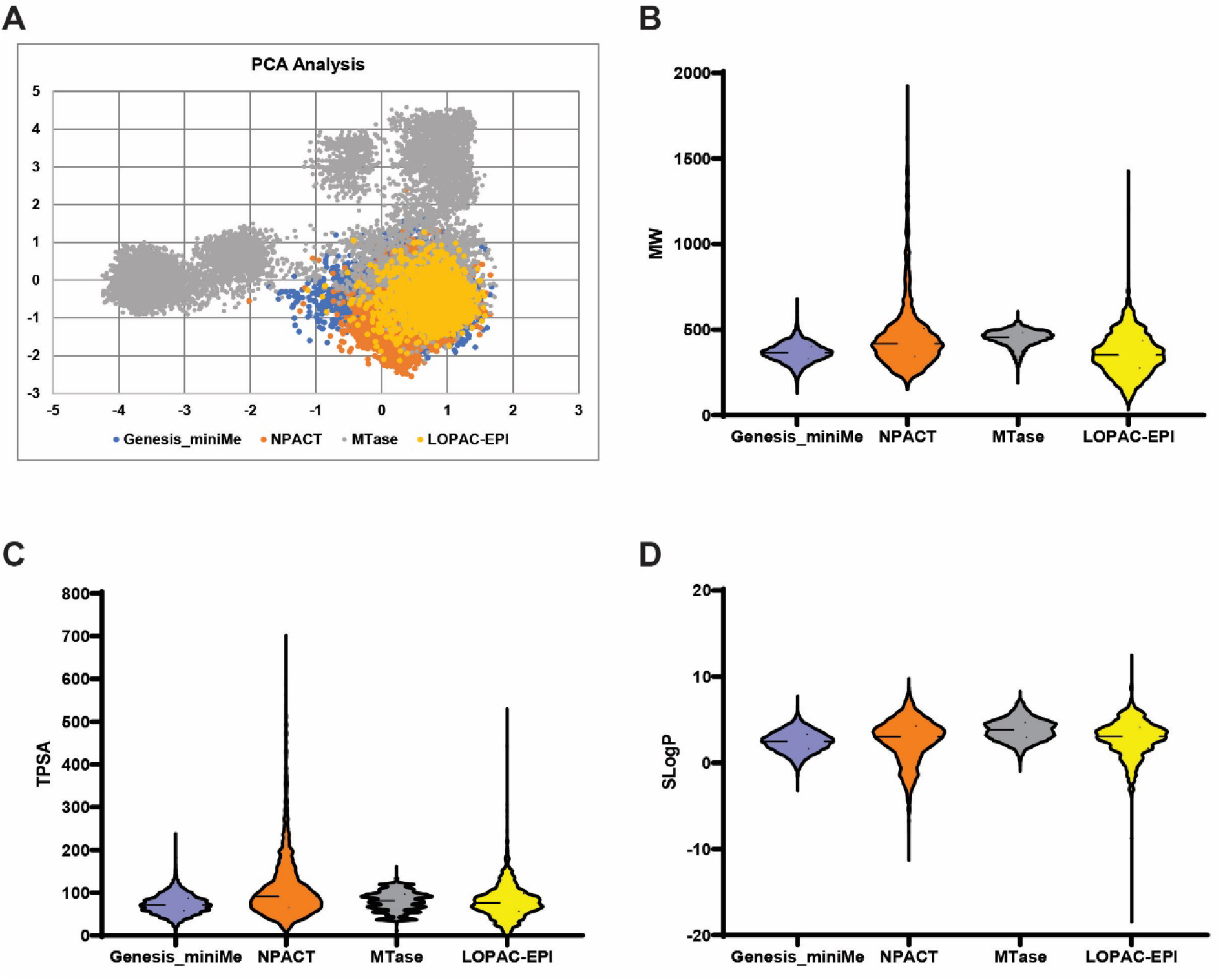
Screening library selection. Four compound libraries were selected to develop the SMMTase target class profile. A) Principal component analysis describing the overall chemical diversity of the annotated libraries (NPACT and LOPAC-Epi) and the chemical diversity libraries (Genesis and MTase). B) Molecular weight distribution of compounds in each compound set. C) Total polar surface area distribution in each library. D) Hydrophobicity plot of all libraries.

### Primary Screen and Assay Performance

The six MTase targets NNMT, HNMT, COMT, PNMT, GNMT, and GAMT were screened in 1536-well plates using the established assay protocols against 27,574 compounds at a single concentration of 40 μM. Compounds which showed at least 50% inhibition against at least one SMMTase were selected for follow-up testing in 11-point dose response against all six MTase assays plus the counter assay (Figure 4A). In this way, all hit compounds were tested twice (HTS at single concentration and cherry-pick follow-up) across the entire assay set. The Z’ for all assay plates was between 0.8 and 0.9 with few exceptions (Figure 4B). Signal-to-background values (positive signal control: full biochemical reaction, negative signal control: no addition of substrate were typically between 4 and 10, with variation in S:B outside that range not affecting Z’ assay quality (Figure 4C). All assay QC values indicate a robust and well-performing set of assays according to standard HTS practices^43^.

**Figure 4.**
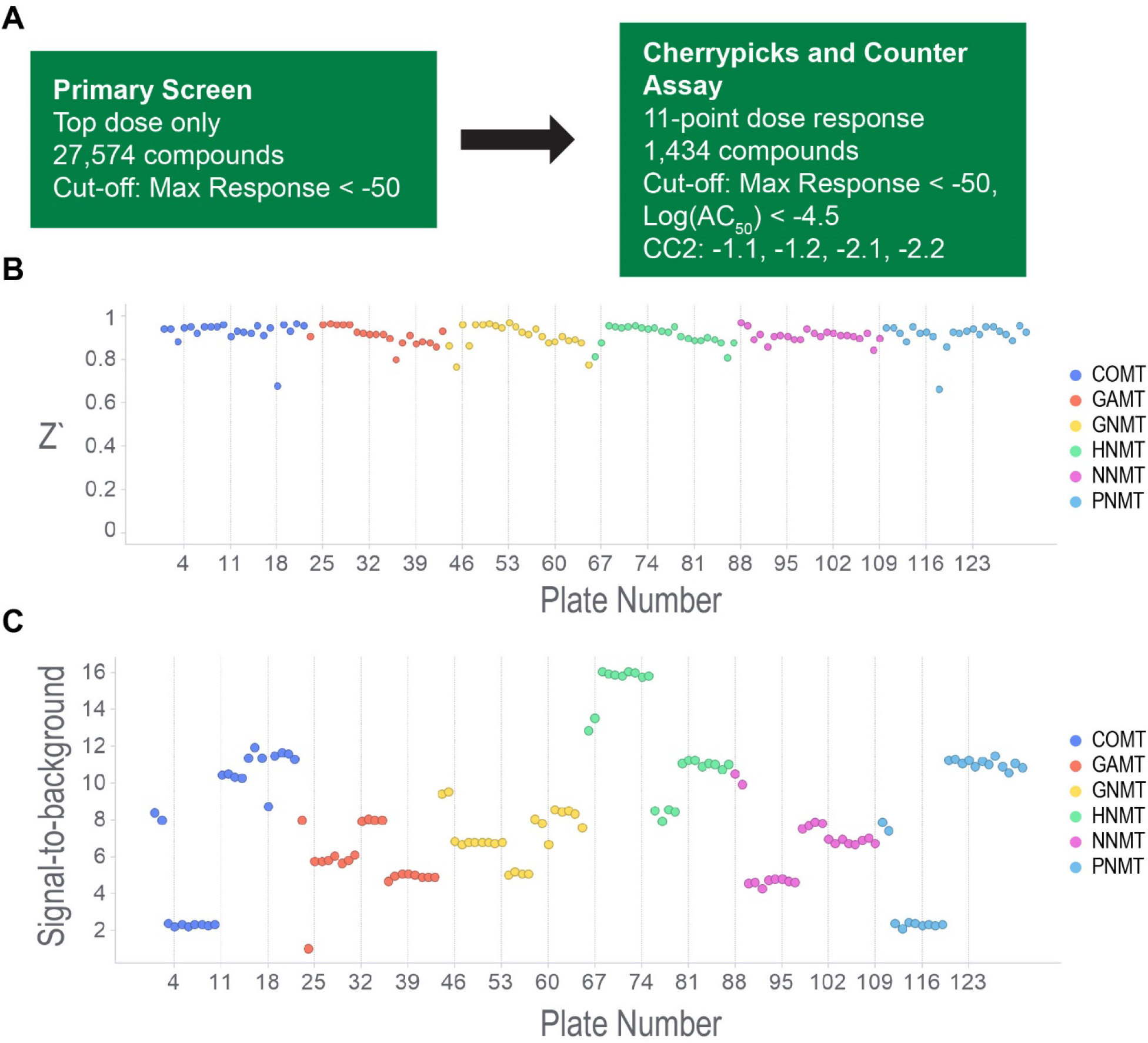
High-throughput screening overview and assay performance. A) A total of 27,574 compounds were screened against NNMT, HNMT, COMT, PNMT, GNMT, and GAMT for MTase inhibition. The initial screen was performed at a concentration of 40 μM, and hits were scored based on a maximum response cut-off of −50, indicating >50% inhibition. 1,434 compounds met this criteria and were then assayed against each MTase and counter-assay in 11-point dose response. Final hits were selected on the basis of maximum response (< −50), Log(AC50) < −4.5, and curve class (−1.1, −1.2, −2.1, −2.2 – see Methods for explanation of curve class categories). B) Z’ for all 1536-well assay plates in the primary screen, calculated from 32 positive- and negative-control wells on each plate. C) Signal-to-background (S/B) for all plates in the primary screen calculated from 32 positive- and negative-control wells on each plate.

In the primary screen of 27,574 compounds, a hit was defined as any compound with at least 50% inhibition. HNMT had the highest initial hit rate of 511 compounds (1.82%) while GAMT had the lowest at 102 compounds (0.36%) (Figure 5B). The rank order for initial hits was HNMT (1.82%) > NNMT (1.44%) > GNMT (1.08%) > COMT (1.07%) > PNMT (0.82%) > GAMT (0.36%). From these hits across all six HTS assays we identified 1,434 unique compounds, which were selected for confirmation- and counter-screening. Activity confirmation for these 1,434 cherrypicked compounds was performed in 11-point dose response (top dose 40 μM, 1:3 serial dilutions) against COMT, HNMT, NNMT, GNMT, PNMT, GAMT, and a counter-assay which used 1 μM SAH in place of enzyme. From these dose response curves we then separated active compounds into two groups: high quality actives and low-quality actives. High quality actives are compounds with complete dose-response curves, a maximum inhibition of over 50%, and IC_50_ < 10 μM. Low quality actives produced incomplete curves or single point activity, but their inclusion in the overall analysis could lend insight into chemical features important for selection/exclusion of members of this target class.

**Figure 5.**
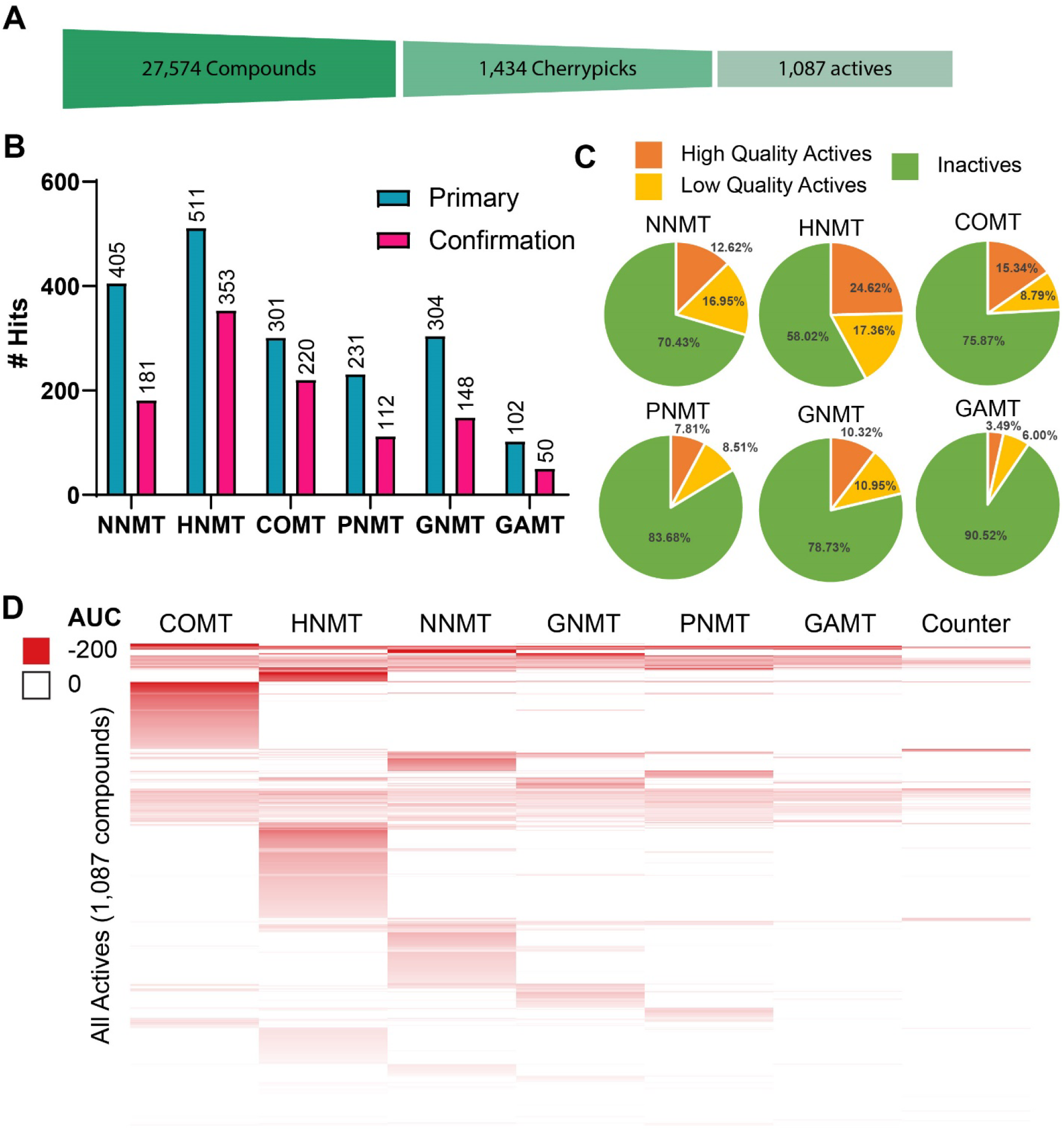
Hit rates for cherrypicked compounds. A) Cartoon depiction of the funnel-down approach to identifying hits. A total of 28,144 compounds were screened and 1,434 compounds were identified as initial hits suitable for cherrypicking. Out of those initial hits a total of 1,087 unique compounds were confirmed across each MTase assay. B) Initial hits for each SMMTase (turquoise) and confirmed hits (magenta). Note that hit numbers do not account for specificity, which is reflected in D. C) 1,434 compounds from the primary screen were selected for cherrypicking. Each compound was assayed against each Mtase and a counter assay in 11-point dose response. Hits were the identified as high-quality actives (complete dose-response curves, max Response ≤ −50, and IC50 ≤ 10 μM), low quality actives (shallow curves, single-point activity, or inconclusive), or inactive. D) Heat map showing activity profile of all active compounds against COMT, HNMT, NNMT, GNMT, PNMT, GAMT, and the counter assay. Compounds in this heat map are clustered by total activity across all seven assays.

Out of the 1,434 cherrypicked compounds, a total of 1,087 unique compounds had their activity reconfirmed in the dose response format (Figure 5B). 54 compounds were active in the counter assay (3.7 %), 8 with IC_50_ values below 10 μM, and were excluded from further analyses. 293 cherrypicked compounds (20.3%) failed to reconfirm. Again, HNMT had the greatest number of high-quality active compounds (353/1,434 compounds, 24.62%) for a final hit rate of 353/28,144 (1.25%). GAMT had the lowest number (50/1,434 high-quality actives, 3.49%) for a final hit rate of 50/28,144 compounds, or 0.18%. The confirmed hit rate (relative to full compound set of 27,574 molecules) rank order shifted to HNMT (1.25%) > COMT (0.78%) > NNMT (0.64%) > GNMT (0.53%) > PNMT (0.40%) > GAMT (0.18%).

To consider the broader-scale perspective on these hit compounds we collapsed each dose response curve to a single area-under-the-curve (AUC) value and generated a heat map summary of the general activity data (Figure 5D). Lower AUC values indicate more potent and efficacious inhibitors of a given enzyme. From this heat map one can quickly survey which enzymes had multiple specific hits. For example, COMT had 220 total hits and 148 of these compounds were specific to COMT. HNMT (219/353 specific hits) and NNMT (92/181 specific hits) also had high specific hit rates. Conversely enzymes like GNMT (51/148 specific hits), PNMT (31/112 specific hits), and GAMT (1/50 specific hits) had low specific hit rates. These data could be used to inform future target screening campaigns. One could draw conclusions such as HNMT, COMT, and NNMT are the most “druggable” targets based on the high overall hit rates and high specificity hit rates. GNMT, PNMT, and GAMT, on the other hand, present more challenging targets for drug development because of their low overall and specific hit rates. These hit rate and specificity analyses make up one angle of these SMMTase target class profiles.

### Chemical clustering analysis

To generate more detailed target class profiles of our SMMTases we grouped compounds by chemical similarity using Tanimoto coefficients as a metric for chemical relatedness. We then plotted the activity of each compound against each SMMTase as AUC values and generated a heat map to survey structureactivity relationships (Figure 6A). A block of compounds compounds active against a single SMMTase indicates a chemotype suitable for a potential starting point for probe development. Another powerful use of this structural clustering analysis is the ability to identify non-specific pan-inhibitors of a target class. A ‘pan-active’ hit can be leveraged as high-quality tool molecules for studying a target class.

**Figure 6.**
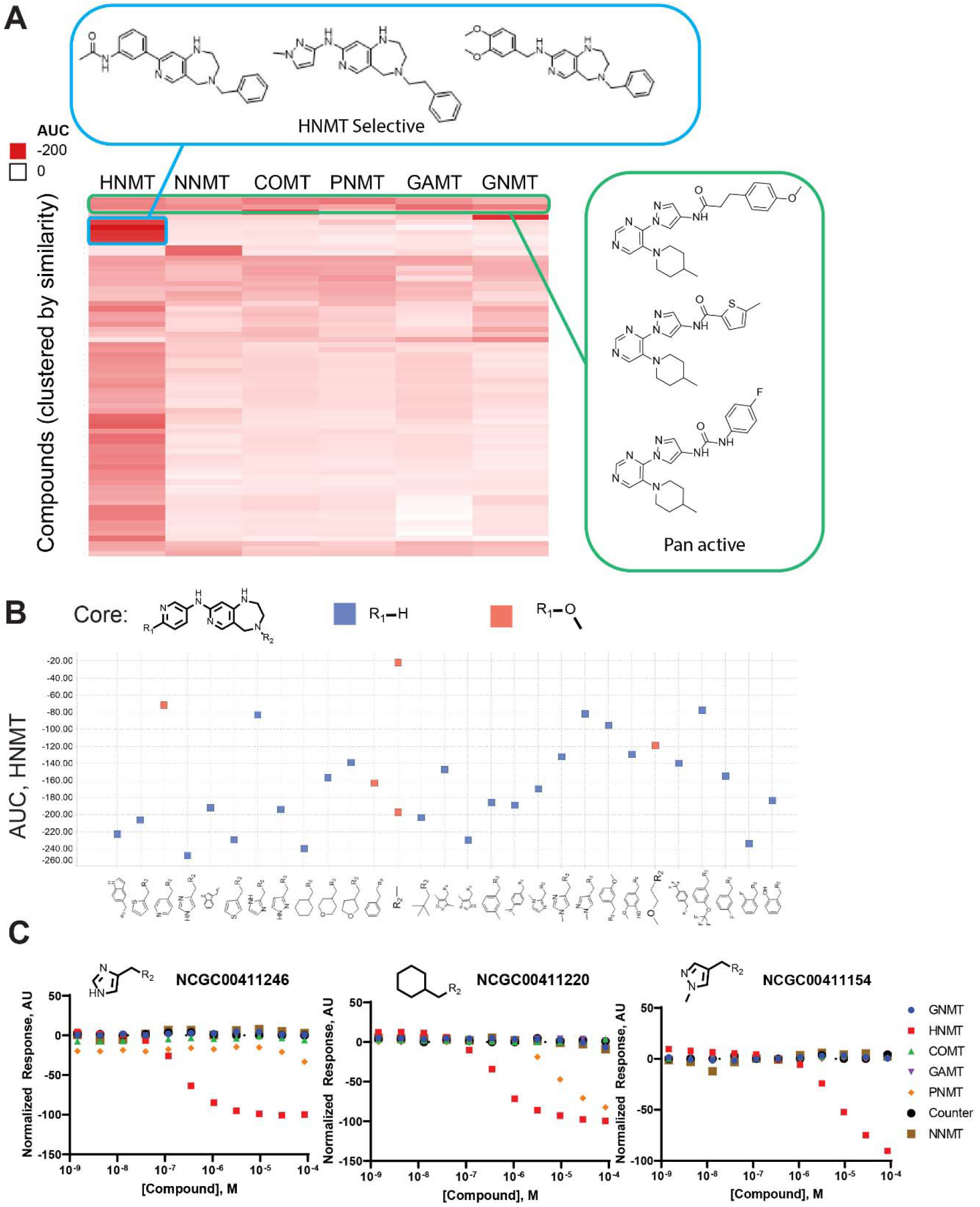
**A)** Cluster analysis of Diversity Library hits. Compounds were clustered according to chemical similarity (based on Tanimoto coefficient) and a heat map was generated to show relative activity. Chemical clusters with high activity, such as the HNMT selective cluster (blue) reveal selective chemotypes for a single SMMTase. Bands of high activity indicate pan inhibitors of the representative SMMTases (green). B) Chemotype of top HNMT-selective hit enables cheminformatic R-group decompositions. R group decomposition was performed using TIBCO Spotfire (TIBCO Software Inc.). Compound activity (displayed as area under the curve, AUC) was plotted as a function of R_2_ functional group identity. R_1_ group identity is color coded blue for hydrogen and orange for methoxy. C) The R2 group in NCGC00411246 yields a highly active and specific compounds. R groups which result in reduced specificity (e.g., NCGC00411220) or reduce activity (e.g., NCGC00411154) we also observed.

### HNMT-specific compounds as a launching pad for drug discovery

One of the ways target class profiling can be leveraged is as a launchpad for more in-depth, target-focused drug development. HNMT was selected as an example because of the relatively high general and specific hit rates in our collections (Figure 5B). We identified a chemotype of HNMT-selective compounds, which all contained a 2,3,4,5-tetrahydro-1H-pyrido[4,3-e][1,4]diazepine core (Figure 6B). These compounds were all from the Genesis MiniMe library (9,856 compounds), which is a subsampling of our larger Genesis collection (>100,000 compounds). We did further SAR expansions using the 2,3,4,5-tetrahydro-1H-pyrido[4,3-e][1,4]diazepine core as a substrate query to search against our NCATS screening collection and identified 254 additional compounds which were not part of the initial screening, set. Each of these compounds was tested in 11-point dose response against HNMT. All hit compounds were tested for selectivity against all SMMTase assays and counter-screened for assay interference. We identified a sub-chemotype of the original diazepine core containing 22 HNMT-inhibitors (Figure 6B). R-group decomposition allowed us to perform preliminary structure-activity analysis to investigate the influence of functional groups on activity and specificity. For example, our best hit contained a methyl-imidazole group at the R2 position (Figure 6C). Introduction of a phenyl ring at R2 not only results in lower activity, but also loss of specificity for HNMT.

To further investigate the SAR of these diazepine compounds we also used molecular modeling of our representative screening hit NCGC00411044 (Figure 7). Docking was performed using an X-ray crystal structure of HNMT (PDB code: 1JQD) bound to the SAM cofactor and the substrate histamine (Figure 7A). Based on the crystal structure, the adenine core of SAM forms H-bond interaction with Ser120, the hydroxyls on the sugar part form multiple H-bonds with Gly60 Glu89, and Gln94. The terminal-NH_2_ group also forms H-bond interactions with the Gly60 and Ile142 backbone carbonyl oxygen. The docking model of NCGC00411044 revealed that the pyridine nitrogen maintains the key interaction with Ser120, which mimic the interaction mode of SAM, and the diazepine core forms two H-bonds with the surrounding residues Gly60 and Gln143 (Figure 7B-D). In addition, the thiophene moiety is positioned into a relatively hydrophobic pocket with pi-stacking to several residues with aromatic side chains including Tyr15, Phe19, Phe22, Gln143, Tyr146 and Tyr147.

**Figure 7.**
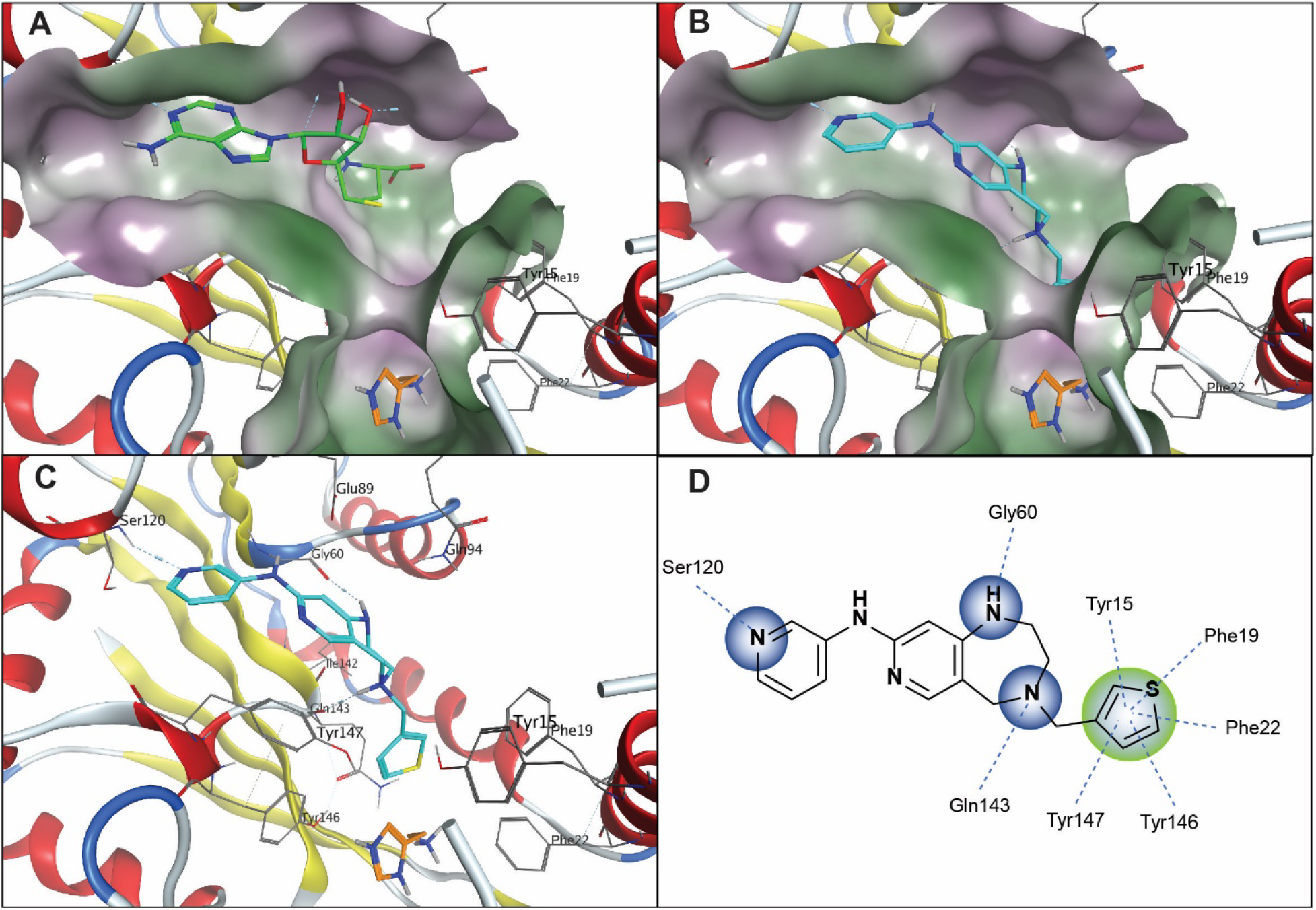
A) Co-crystal structure of HNMT (PDB ID: 1JQD) cofactor SAM is showing in green stick mode and the substrate histamine is showing in orange stick mode. B) and C) Docking model of the screening hit NCGC00411044 at SAM binding pocket. NCGC00411044 is showing in cyan stick mode. D) 2D ligand-protein interaction map to show the specific H-bond interaction and pi-stacking interactions. The molecular surface of the binding site is depicted in hydrophobicity contour (hydrophilic = purple; lipophilic=green), and the dotted lines indicate H-bond interactions. Figures are prepared using Molecular Operating Environment (MOE) computational software.

This docking hypothesis was corroborated by the SAR expansion data set (Table S1). In brief, with the highly conserved diazepine core, compounds with both pyridine and terminal aromatic ring on the other side (NCGC00411145, NCGC00411246, NCGC00411044, and NCGC00411168) showed best activity as they can interact with Ser120 and form potential multiple pi-stacking with the aromatic side chains. From the binding model, the position of the pyridine ring nitrogen also impacts compound activity, missing the nitrogen (NCGC00411199, NCGC00411050 and NCGC00411115) or nitrogen at the *para*-position (NCGC00411201 and NCGC00411196) cause at least 10-fold potency loss. On the other hand, the aromatic thiophene in NCGC00411044 forms strong pi-pi interactions with the surrounding residues, so analogs missing this aromatic moiety (NCGC00411147, NCGC00411078, and NCGC00411229) also resulted in potency loss. Finally, if the compounds are completely lacking these two types of critical interactions, they showed no activity in the biochemical assays. These findings are highly consistent with the docking hypothesis.

### Pan-active inhibitors are putative SMMTase-selective

In addition to the HNMT-selective compounds discussed above, we also identified a chemotype of non-specific compounds, which demonstrate no signs of assay interference (Figure 6A). We rationalized that if members of this chemotype showed SMMTase-specific activity, and not broader MTase activity, it could represent a novel class of biological probe molecule. Tool molecules are staples of kinase biology and can be used to distinguish between kinase-dependent and kinase-independent pathways^44^ and to identify kinases in phosphorylation cascades^45^. Probe molecules in this sense are not available for MTases. To this end, we selected two members of the putative pan-SMMTase inhibitor chemotype, NCGC00519772 and NCGC00519779. Both compounds inhibit PNMT, HNMT, NNMT, GNMT, GAMT, and COMT at high micromolar concentrations. To determine if these compounds were selective for SMMTases or general inhibitors of the larger MTase family we submitted two such compounds for testing against a panel of 39 protein and DNA methyltransferases (Reaction Biology, Malvern, PA). Of the 39 additional MTases, only four (PRMT3, PRMT7, PRMT8, and SMYD7) were weakly inhibited by the compounds (Figure 8). Based on these data, NCGC00519772 and NCGC00579779 are putatively selective for SMMTases.

**Figure 8.**
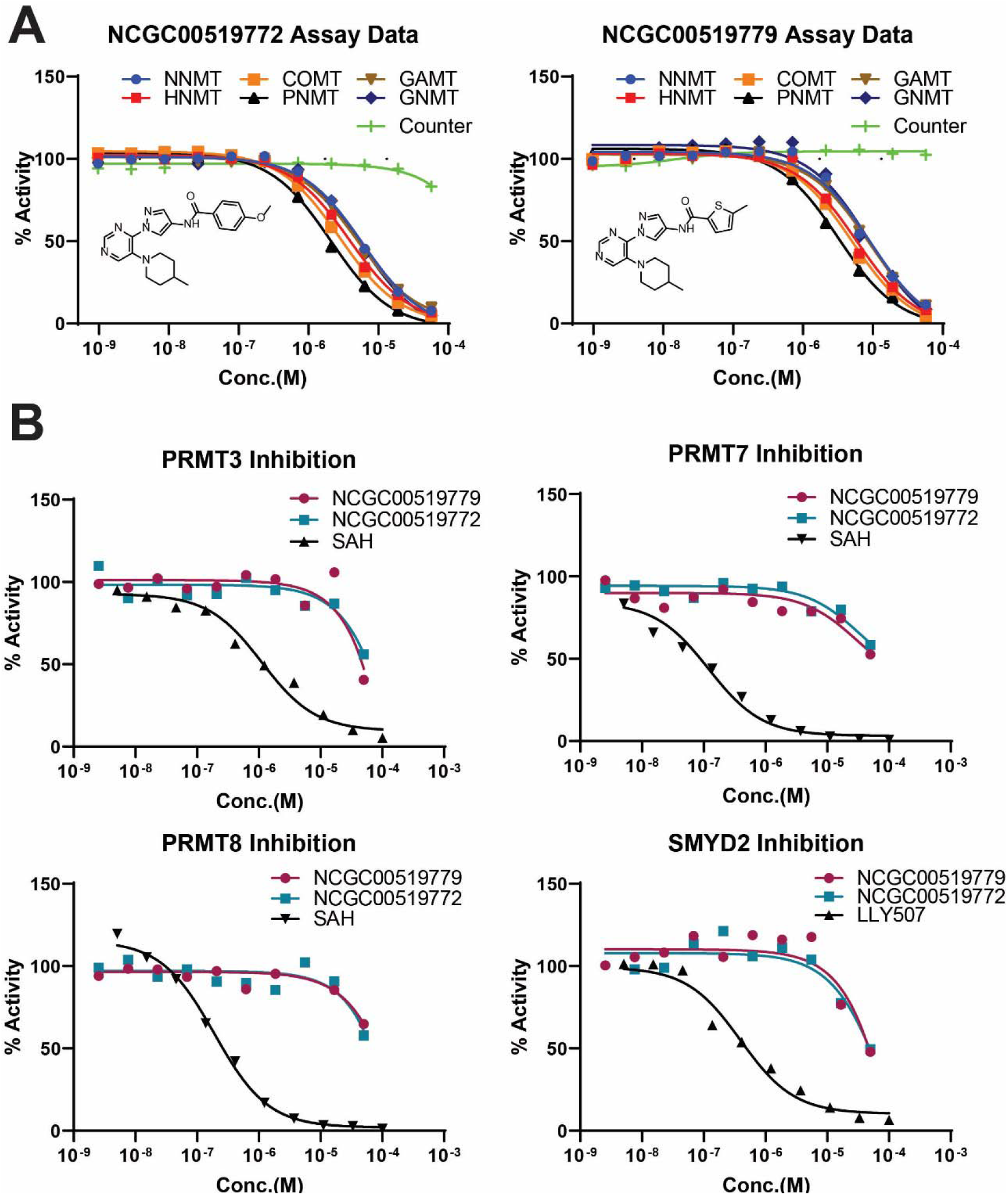
Pan inhibitors of SMMTase activity are selective for SMMTase subfamily. A) NCGC00519779 and NCGC00519772 were activite against the SMMTases PNMT, COMT, HNMT, GNMT, NNMT, and GAMT. No assay interference was observed in the counter-assay. B) NCGC00519779 and NCGC00519772 were submitted to Reaction Biology (Malvern, PA) for additional testing against 39 protein and DNA MTases. Each compound was tested in 10-point dose response, 1:3 serial dilution, 50 μM highest concentration. A reference inhibitor was also used for each MTase (black curves). Only PRMT3, PRMT7, PRMT8, and SMYD2 had observable activity, falling under the “low quality active” category (incomplete dose-response curve; activity at a single dose).

The assay development, compound libraries, and informatics capabilities described herein constitute a platform for target class profiling of SMMTases. By screening four libraries (totaling 27,574 compounds) across the SMMTases PNMT, HNMT, GNMT, NNMT, GAMT, and COMT we created activity profiles for this target class, which can be leveraged for development of tool molecules or targeted screening. From these data we identified the putative SMMTase subclass-selective probes NCGC00519772 and NCGC00519779. At current writing these probes are being pursued to understand the mechanism of selective and further develop the chemotype. We presented HNMT as an example of leveraging these target class profiles to launch a targeted probe development campaign. At NCATS we are applying this framework to other SMMTase candidates with the added goal of developing cellular assays to create cell-active probes. There are also many exciting opportunities for expanding on this work by adding other SMMTases to the portfolio.

## Methods

### Reagents

Recombinant human PNMT, NNMT, HNMT, COMT, GNMT, and GAMT were all acquired from Novus Biologicals (Centennial, CO). MTase Glo Kits, including the SAM cofactor we all purchased from Promega (Madison, WI). Norepinephrine, nicotinamide, glycine, guanidinoacetate, and histamine were all purchased from Sigma Aldrich (St. Louis, MO). Initial assay development was performed in 384-well Lumitrac plates (White, flat bottom, medium binding, Cat# 781075, Greiner Bio-One, Monroe, NC). HTS assays were done in Aurora 1536 plates (white, flat bottom, medium binding Cat# AWK010000A, Aurora Microplates, Scottsdale, AZ). All data were analyzed and processed using in-house tools and TIBCO Spotfire 6.0.0 (Tibco Software Inc., Cambridge, MA. https://spotfire.tibco.com/). Data visualizations were generated in GraphPad Prism (GraphPad Software, San Diego, CA), TIBCO Spotfire, and Adobe Illustrator (Adobe, San Jose, CA).

### Determining HTS Assay Parameters

K_M_ values of each substrate for their respective enzyme were determined using the method of initial rates. Briefly, MTase enzymes were diluted to 5-30 nM in 20 mM Tris (pH 8.0), 150 mM NaCl, 12 mM MgCl2, 4 mM EDTA, 0.1% BSA, 100 μM SAM. Reactions were initiated by adding substrate at various concentrations. Reactions were allowed to proceed for discreet time points and progress was terminated by adding the MTase Glo reagent to deplete excess SAM. Reaction progress was measured as a function of SAM to SAH conversion using the MTase Glo Kit (Promega). K_M_ values for SAM were determined by the same method but holding substrate concentrations constant at 100 μM and varying the concentration of SAM. Initial rates were determined by plotting the generation of SAH as a function of time to generate a linear slop. These rates were then plotted as a function of substrate concentration and fit to the Michaelis-Menten Equation to determine K_M_. The only exception is for PNMT, which was fit to a substrate inhibition curve.

HTS Assay duration was determined by running a time course experiment using the previously determined K_M_ values for substrate and SAM. For each enzyme a mixture of SAM (at the determined K_M_ concentration) and substrate (at the determined K_M_ concentration) were prepared in 20 mM Tris (pH 8.0), 150 mM NaCl, 12 mM MgCl_2_, 4 mM EDTA, 0.1% BSA. Reactions were initiated by adding SMMTase to a final concentration of 7.5, 15, or 30 μM. Reaction progress was recorded at regular intervals by quenching the reaction with the addition of MTase Glo reagent to consume excess SAM. The 20% conversation point was identified by comparing reaction progress to a SAH standard curve. Ideal reaction duration and enzyme concentration were selected based on the combination which achieved 10-20% conversion in 20-30 minutes.

### Primary screen and data analysis

NNMT, COMT, HNMT, PNMT, GNMT, and GAMT were all subject to primary screening against a panel of 27,574 compounds. Primary screening was performed against single dose of compound (40 μM final) in 1536-well format using automated HTS protocols (Table 1). SAM and substrate concentrations used were based on the determined K_M_ values (Table 2). Detailed assay protocols for each SMMTase can be found in Supplemental Methods. To determine compound activity in the primary assay, raw plate reads for each compound were first normalized relative to the positive control (sinefungin, −100% activity, full inhibition) and DMSO only wells (basal, 0% activity) using in-house informatics tools.

### Cherrypicks, counter-screening, and hit selection

Compounds which showed at least 50% inhibition in any of the six assays in the primary screen were selected for cherrypicking. Cherrypicked compounds were replated using fresh stock in dose response (Top dose 40 μM final, 11-points, 1:3 dilution series) and manually run against all six SMMTase assays (Table 1) and a counter assay. The concentration-response data for each sample was plotted and modeled by a four parameter logistic fit yielding IC_50_ and efficacy (maximal response) values as previously described. Compounds were designated as Class 1–4 according to the type of concentration–response curve (CRC) observed. In brief, Class −1.1 and −1.2 were the highest-confidence complete CRCs containing upper and lower asymptotes with efficacies ≥ 80% and < 80%, respectively. Class −2.1 and −2.2 were incomplete CRCs having only one asymptote with efficacy ≥ 80% and < 80%, respectively. Class −3 CRCs showed activity at only the highest concentration or were poorly fit. Class 4 CRCs were inactive having a curve-fit of insufficient efficacy or lacking a fit altogether. Compounds which showed any activity in the counter assay were excluded from hit selection and further analysis. Hits were defined as either high-quality actives (complete dose response curves, max Response ≤ −50, and IC_50_ ≤ 10 μM), low-quality actives (shallow curves, single-point activity, or inconclusive), or inactive (no activity).

### Clustering of compounds by activity outcomes

Compounds were clustered hierarchically using TIBCO Spotfire 6.0.0 (Tibco Software Inc., Cambridge, MA. https://spotfire.tibco.com/) based on their activity outcomes from the primary or follow up screens across different methyltransferases. Compound’s AUC (Area Under the Curve) calculated based on the qHTS data analysis and curve fittings were utilized for clustering. In the heatmap, darker red indicates compounds that are more potent and efficacious, i.e. high-quality actives, and lighter red indicates less potent and efficacious compounds. If a compound didn’t show any activity in an assay, it was highlighted as white in the heatmap.

### Structural clustering analysis and HNMT inhibitor SAR expansion

The cherrypicked compounds identified from qHTS screens against Genesis_miniMe, annotated and MTase chemical libraries were further clustered using MACCS Structural Keys (Bit packed) (FP:BIT_MACCS) fingerprint and Tanimoto coefficient as similarity metric in Molecular Operating Environment software (MOE, https://www.chemcomp.com/), respectively. The similarity and overlap level were both set at 75. The most significant enriched scaffold for HNMT is 2,3,4,5-tetrahydro-1H-pyrido[4,3-e][1,4]diazepine. Members in this cluster showed comparable IC_50_ values in HNMT biochemical inhibition assay and great selectivity against all other five methyltransferases. We further did SAR (Structure-Activity Relationship) expansion using the core 2,3,4,5-tetrahydro-1H-pyrido[4,3-e][1,4]diazepine as the substructure query to search against the entire NCATS screening collection, and identified 254 compounds sharing the same scaffold. These 254 SAR expansion compounds were then plated in 11-pt dose response using freshly prepared stock and tested in the follow-up assays.

### Molecular modeling

HNMT co-crystal structure (PDB code 1JQD) was prepared for the docking studies using MOE molecular modeling software. QuickPrep module was used to set the correct protonation state for protein, and water molecules farther than 4.5 Å from the active site was deleted. All hydrogens were added to the protein and partial charges were attributed to the protein atoms using Amber99 force field. In MOE docking, ligand conformations were generated on-the-fly, and the active site was defined by glycine-rich loop and the hinge region residues without pharmacophore constraints. The first round of docking to roughly filter out unfavorable binding poses used London dG scoring function, then the top 30 ranking poses were further refined using GBVI/WSA dG scoring function. Finally, the top 5 ranking docking poses after refinement were selected and manually inspected based on the binding orientations and specific interactions.

### Expanded methyltransferase profiling

NCGC00519779, NCGC00519772, NCGC00411145, and NCGC00411044 were assessed for inhibition against a panel of recombinant human protein methyltransferases by commercial services from Reaction Biology Corp (Malvern, PA). Reaction Biology Corp tested each compound in a 10-point dose response with a 1:3 dilution series starting at 50 μM against 39 protein methyltransferases.

## Supporting information

Supporting information is available free of charge at _.

Supporting Methods. Detailed assays information for each SMMTase used in this study.

## Author Contribution

Q.M.H. developed and designed experiments. Q.M.H. and I.C. conducted experiments. Q.M.H., M.S., and H.G. conducted data analysis. M.S. conducted all modelling experiments. Q.M.H., M.S., and M.D.H. wrote and edited the manuscript.

## Notes

The authors declare no competing financial interests.

## Acknowledgements

We would like to thank the compound management group at the National Center for Advancing Translational Science for preparing all compound plates. The work was supported by the Intramural Research Program of the National Center for Advancing Translational Science, National Institutes of Health. Q.M.H. was funded by Eli Lilly.

## References

1. Barnash, K. D.; James, L. I.; Frye, S. V., Target class drug discovery. Nat Chem Biol 2017, 13 (10), 1053–1056.

2. Copeland, R. A.; Solomon, M. E.; Richon, V. M., Protein methyltransferases as a target class for drug discovery. Nat Rev Drug Discov 2009, 8 (9), 724–32.

3. Heilker, R.; Wolff, M.; Tautermann, C. S.; Bieler, M., G-protein-coupled receptor-focused drug discovery using a target class platform approach. Drug Discov Today 2009, 14 (5-6), 231–40.

4. Carter, A. J.; Kraemer, O.; Zwick, M.; Mueller-Fahrnow, A.; Arrowsmith, C. H.; Edwards, A. M., Target 2035: probing the human proteome. Drug Discov Today 2019, 24 (11), 2111–2115.

5. Posy, S. L.; Hermsmeier, M. A.; Vaccaro, W.; Ott, K. H.; Todderud, G.; Lippy, J. S.; Trainor, G. L.; Loughney, D. A.; Johnson, S. R., Trends in kinase selectivity: insights for target class-focused library screening. J Med Chem 2011, 54 (1), 54–66.

6. Benn, C. L.; Dawson, L. A., Clinically Precedented Protein Kinases: Rationale for Their Use in Neurodegenerative Disease. Front Aging Neurosci 2020, 12, 242.

7. Goldstein, D. M.; Gray, N. S.; Zarrinkar, P. P., High-throughput kinase profiling as a platform for drug discovery. Nat Rev Drug Discov 2008, 7 (5), 391–7.

8. Glukhova, A.; Draper-Joyce, C. J.; Sunahara, R. K.; Christopoulos, A.; Wootten, D.; Sexton, P. M., Rules of Engagement: GPCRs and G Proteins. ACS Pharmacol Transl Sci 2018, 1 (2), 73–83.

9. Andrews, S. P.; Brown, G. A.; Christopher, J. A., Structure-based and fragment-based GPCR drug discovery. ChemMedChem 2014, 9 (2), 256–75.

10. Hamamoto, R.; Nakamura, Y., Dysregulation of protein methyltransferases in human cancer: An emerging target class for anticancer therapy. Cancer Sci 2016, 107 (4), 377–84.

11. Ferreira de Freitas, R.; Ivanochko, D.; Schapira, M., Methyltransferase Inhibitors: Competing with, or Exploiting the Bound Cofactor. Molecules 2019, 24 (24).

12. Liscombe, D. K.; Louie, G. V.; Noel, J. P., Architectures, mechanisms and molecular evolution of natural product methyltransferases. Nat Prod Rep 2012, 29 (10), 1238–50.

13. Lenz, T.; Poot, P.; Weinhold, E.; Dreger, M., Profiling of methyltransferases and other S-Adenosyl-L-homocysteine-binding proteins by Capture Compound mass spectrometry. Methods Mol Biol 2012, 803, 97–125.

14. Horning, B. D.; Suciu, R. M.; Ghadiri, D. A.; Ulanovskaya, O. A.; Matthews, M. L.; Lum, K. M.; Backus, K. M.; Brown, S. J.; Rosen, H.; Cravatt, B. F., Chemical Proteomic Profiling of Human Methyltransferases. J Am Chem Soc 2016, 138 (40), 13335–13343.

15. Kozbial, P. Z.; Mushegian, A. R., Natural history of S-adenosylmethionine-binding proteins. BMC Struct Biol 2005, 5, 19.

16. Wlodarski, T.; Kutner, J.; Towpik, J.; Knizewski, L.; Rychlewski, L.; Kudlicki, A.; Rowicka, M.; Dziembowski, A.; Ginalski, K., Comprehensive structural and substrate specificity classification of the Saccharomyces cerevisiae methyltransferome. PLoS One 2011, 6 (8), e23168.

17. Petrossian, T. C.; Clarke, S. G., Uncovering the human methyltransferasome. Mol Cell Proteomics 2011, 10 (1), M110 000976.

18. Schubert, H. L.; Blumenthal, R. M.; Cheng, X., Many paths to methyltransfer: a chronicle of convergence. Trends Biochem Sci 2003, 28 (6), 329–35.

19. Fenwick, M. K.; Ealick, S. E., Towards the structural characterization of the human methyltransferome. Curr Opin Struct Biol 2018, 53, 12–21.

20. Bastos, P.; Gomes, T.; Ribeiro, L., Catechol-O-Methyltransferase (COMT): An Update on Its Role in Cancer, Neurological and Cardiovascular Diseases. Rev Physiol Biochem Pharmacol 2017, 173, 1–39.

21. Jarrold, J.; Davies, C. C., PRMTs and Arginine Methylation: Cancer’s Best-Kept Secret? Trends Mol Med 2019, 25 (11), 993–1009.

22. Meng, H.; Cao, Y.; Qin, J.; Song, X.; Zhang, Q.; Shi, Y.; Cao, L., DNA methylation, its mediators and genome integrity. Int J Biol Sci 2015, 11 (5), 604–17.

23. Scheer, S.; Ackloo, S.; Medina, T. S.; Schapira, M.; Li, F.; Ward, J. A.; Lewis, A. M.; Northrop, J. P.; Richardson, P. L.; Kaniskan, H. U.; Shen, Y.; Liu, J.; Smil, D.; McLeod, D.; Zepeda-Velazquez, C. A.; Luo, M.; Jin, J.; Barsyte-Lovejoy, D.; Huber, K. V. M.; De Carvalho, D. D.; Vedadi, M.; Zaph, C.; Brown, P. J.; Arrowsmith, C. H., A chemical biology toolbox to study protein methyltransferases and epigenetic signaling. Nat Commun 2019, 10 (1), 19.

24. Ma, Z.; Liu, H.; Wu, B., Structure-based drug design of catechol-O-methyltransferase inhibitors for CNS disorders. Br J Clin Pharmacol 2014, 77 (3), 410–20.

25. Akhtar, M. J.; Yar, M. S.; Grover, G.; Nath, R., Neurological and psychiatric management using COMT inhibitors: A review. Bioorg Chem 2020, 94, 103418.

26. Barrow, J. C., Inhibitors of Catechol-O-Methyltransferase. CNS Neurol Disord Drug Targets 2012, 11 (3), 324–32.

27. Robinson, R. G.; Smith, S. M.; Wolkenberg, S. E.; Kandebo, M.; Yao, L.; Gibson, C. R.; Harrison, S. T.; Polsky-Fisher, S.; Barrow, J. C.; Manley, P. J.; Mulhearn, J. J.; Nanda, K. K.; Schubert, J. W.; Trotter, B. W.; Zhao, Z.; Sanders, J. M.; Smith, R. F.; McLoughlin, D.; Sharma, S.; Hall, D. L.; Walker, T. L.; Kershner, J. L.; Bhandari, N.; Hutson, P. H.; Sachs, N. A., Characterization of non-nitrocatechol pan and isoform specific catechol-O-methyltransferase inhibitors and substrates. ACS Chem Neurosci 2012, 3 (2), 129–40.

28. Drinkwater, N.; Gee, C. L.; Puri, M.; Criscione, K. R.; McLeish, M. J.; Grunewald, G. L.; Martin, J. L., Molecular recognition of physiological substrate noradrenaline by the adrenaline-synthesizing enzyme PNMT and factors influencing its methyltransferase activity. Biochem J 2009, 422 (3), 463–71.

29. Pendleton, R. G.; Kaiser, C.; Gessner, G., Studies on adrenal phenylethanolamine N-methyltransferase (PNMT) with S K & F 64139, a selective inhibitor. J Pharmacol Exp Ther 1976, 197 (3), 623–32.

30. Rucci, F.; Cigoli, M. S.; Marini, V.; Fucile, C.; Mattioli, F.; Robbiano, L.; Cavallari, U.; Scaglione, F.; Perno, C. F.; Penco, S.; Marocchi, A., Combined evaluation of genotype and phenotype of thiopurine S-methyltransferase (TPMT) in the clinical management of patients in chronic therapy with azathioprine. Drug Metabolism and Personalized Therapy 2019, 0 (0).

31. Keuzenkamp-Jansen, C. W.; Leegwater, P. A.; De Abreu, R. A.; Lambooy, M. A.; Bokkerink, J. P.; Trijbels, J. M., Thiopurine methyltransferase: a review and a clinical pilot study. J Chromatogr B Biomed Appl 1996, 678 (1), 15–22.

32. Yen, C.-H.; Lin, Y.-T.; Chen, H.-L.; Chen, S.-Y.; Chen, Y.-M. A., The multi-functional roles of GNMT in toxicology and cancer. 2013, 266 (1), 67–75.

33. Nieman, K. M.; Hartz, C. S.; Szegedi, S. S.; Garrow, T. A.; Sparks, J. D.; Schalinske, K. L., Folate status modulates the induction of hepatic glycine N-methyltransferase and homocysteine metabolism in diabetic rats. Am J Physiol Endocrinol Metab 2006, 291 (6), E1235–42.

34. Hsiao, K.; Zegzouti, H.; Goueli, S. A., Methyltransferase-Glo: a universal, bioluminescent and homogenous assay for monitoring all classes of methyltransferases. Epigenomics 2016, 8 (3), 321–39.

35. Brooks, H. B.; Geeganage, S.; Kahl, S. D.; Montrose, C.; Sittampalam, S.; Smith, M. C.; Weidner, J. R., Basics of Enzymatic Assays for HTS. In Assay Guidance Manual, Markossian, S.; Grossman, A.; Brimacombe, K.; Arkin, M.; Auld, D.; Austin, C. P.; Baell, J.; Chung, T. D. Y.; Coussens, N. P.; Dahlin, J.L.; Devanarayan, V.; Foley, T. L.; Glicksman, M.; Hall, M. D.; Haas, J. V.; Hoare, S. R. J.; Inglese, J.; Iversen, P. W.; Kales, S. C.; Lal-Nag, M.; Li, Z.; McGee, J.; McManus, O.; Riss, T.; Saradjian, P.; Sittampalam, G. S.; Tarselli, M.; Trask, O. J., Jr.; Wang, Y.; Weidner, J. R.; Wildey, M.J.; Wilson, K.; Xia, M.; Xu, X., Eds. Bethesda (MD), 2004.

36. Beaudouin, C.; Haurat, G.; Fraisse, L.; Souppe, J.; Renaud, B., Assay of phenylethanolamine N-methyltransferase activity using high-performance liquid chromatography with ultraviolet absorbance detection. 1993, 613 (1), 51–58.

37. Horton, J. R.; Sawada, K.; Nishibori, M.; Zhang, X.; Cheng, X., Two polymorphic forms of human histamine methyltransferase: structural, thermal, and kinetic comparisons. Structure 2001, 9 (9), 837–49.

38. Zhang, J.; Klinman, J. P., Convergent Mechanistic Features between the Structurally Diverse N- and O-Methyltransferases: Glycine N-Methyltransferase and Catechol O-Methyltransferase. J Am Chem Soc 2016, 138 (29), 9158–65.

39. Loring, H. S.; Thompson, P. R., Kinetic Mechanism of Nicotinamide N-Methyltransferase. Biochemistry 2018, 57 (38), 5524–5532.

40. Lotta, T.; Vidgren, J.; Tilgmann, C.; Ulmanen, I.; Melen, K.; Julkunen, I.; Taskinen, J., Kinetics of human soluble and membrane-bound catechol O-methyltransferase: a revised mechanism and description of the thermolabile variant of the enzyme. Biochemistry 1995, 34 (13), 4202–10.

41. Ogawa, H.; Gomi, T.; Takata, Y.; Date, T.; Fujioka, M., Recombinant expression of rat glycine N-methyltransferase and evidence for contribution of N-terminal acetylation to co-operative binding of S-adenosylmethionine. Biochem J 1997, 327 (Pt 2), 407–12.

42. Lipinski, C. A.; Lombardo, F.; Dominy, B. W.; Feeney, P. J., Experimental and computational approaches to estimate solubility and permeability in drug discovery and development settings. Adv Drug Deliv Rev 2001, 46 (1-3), 3–26.

43. Iversen, P. W.; Beck, B.; Chen, Y. F.; Dere, W.; Devanarayan, V.; Eastwood, B. J.; Farmen, M. W.; Iturria, S. J.; Montrose, C.; Moore, R. A.; Weidner, J. R.; Sittampalam, G. S., HTS Assay Validation. In Assay Guidance Manual, Markossian, S.; Grossman, A.; Brimacombe, K.; Arkin, M.; Auld, D.; Austin, C. P.; Baell, J.; Chung, T. D. Y.; Coussens, N. P.; Dahlin, J. L.; Devanarayan, V.; Foley, T. L.; Glicksman, M.; Hall, M. D.; Haas, J. V.; Hoare, S. R. J.; Inglese, J.; Iversen, P. W.; Kales, S. C.; Lal-Nag, M.; Li, Z.; McGee, J.; McManus, O.; Riss, T.; Saradjian, P.; Sittampalam, G. S.; Tarselli, M.; Trask, O. J., Jr.; Wang, Y.; Weidner, J. R.; Wildey, M. J.; Wilson, K.; Xia, M.; Xu, X., Eds. Bethesda (MD), 2004.

44. Dranchak, P.; MacArthur, R.; Guha, R.; Zuercher, W. J.; Drewry, D. H.; Auld, D. S.; Inglese, J., Profile of the GSK published protein kinase inhibitor set across ATP-dependent and-independent luciferases: implications for reporter-gene assays. PLoS One 2013, 8 (3), e57888.

45. Watson, N. A.; Cartwright, T. N.; Lawless, C.; Camara-Donoso, M.; Sen, O.; Sako, K.; Hirota, T.; Kimura, H.; Higgins, J. M. G., Kinase inhibition profiles as a tool to identify kinases for specific phosphorylation sites. Nat Commun 2020, 11 (1), 1684.

