## Supplemental Methods for "Target class profiling of small molecule methyltransferases"

The following tables contain detailed assay steps for each small molecule methyltransferase used in this study. All concentrations in the Description column indicate the input concentration. Final assay conditions, including final reagent concentrations, are detailed in the Notes section.

| Protocol table for COMT HTS MTase-Glo Assay | | | |
| --- | --- | --- | --- |
| Step | Parameter | Value | Description |
| 1 | Reagent | 2 µL | COMT (20 nM) in reaction buffer, columns 1-48 |
| 2 | Reagent | 1 µL | SAM (20 µM) in reaction buffer, columns 1-48 |
| 3 | Controls | 23 nL | DMSO in column 4; sinefungin in DMSO (0 µM – 40 µM) in 7-point 1:2 dilution series (n = 2) in column 2 |
| 4 | Library compounds | 23 nL | Columns 5-48 |
| 5 | Reagent | 1 µL | Norepinephrine (60 µM) in reaction buffer, columns 2-48 |
| 6 | Time | 20 min | Incubation |
| 7 | Reagent | 1 µL | MTase-Glo reagent (5X), columns 1-48 |
| 8 | Time | 30 min | Incubation |
| 9 | Reagent | 5 uL | MTase-Glo Detection reagent, columns 1-48 |
| 10 | Time | 30 min | Incubation, luminescence evolution |
| 11 | Detection | Luminescence | ViewLux uHTS Microplate Imager (PerkinElmer) |
| Step | Notes | | |
| 1, 2, 5 | White Medium Binding 1536-well plates (white, flat bottom, medium binding Cat# AWK010000A, Aurora Microplates, Scottsdale, AZ); Reaction buffer: 50 mM Tris, pH 8.0, 3 mM MgCl_2_, 1 mM EDTA, 50 mM NaCl, 1 mM DTT, and 0.1 mg/mL BSA; refer to tables 2 and S1 for specific reagent concentrations | | |
| 3, 4 | Pintool transfer | | |
| 6, 8, 10 | Room temperature | | |
| 6 | Final reaction conditions: 10 nM COMT, 5 µM SAM, 15 µM norepinephrine, 50 mM Tris, pH 8.0, 3 mM MgCl_2_, 1 mM EDTA, 50 mM NaCl, 1 mM DTT, and 0.1 mg/mL BSA | | |
| 8 | Conversion of SAH to ADP; MTase Glo Kit (Promega, Madison, WI) | | |
| 10 | Conversion of ADP to ATP and detection by UltraGlo luciferase; MTase Glo Kit (Promega, Madison, WI) | | |
| 11 | Settings: 20 s exposure, 1X binning, high gain, medium speed | | |

| Protocol table for PNMT HTS MTase-Glo Assay | | | |
| --- | --- | --- | --- |
| Step | Parameter | Value | Description |
| 1 | Reagent | 2 µL | PNMT (40 nM) in reaction buffer, columns 1-48 |
| 2 | Reagent | 1 µL | SAM (20 uM) in reaction buffer, columns 1-48 |
| 3 | Controls | 23 nL | DMSO in column 4; sinefungin in DMSO (0 µM – 40 µM) in 7-point 1:2 dilution series (n = 2) in column 2 |
| 4 | Library compounds | 23 nL | Columns 5-48 |
| 5 | Reagent | 1 µL | Norepinephrine (60 µM) in reaction buffer, columns 2-48 |
| 6 | Time | 30 min | Incubation |
| 7 | Reagent | 1 µL | MTase-Glo reagent (5X), columns 1-48 |
| 8 | Time | 30 min | Incubation |
| 9 | Reagent | 5 uL | MTase-Glo Detection reagent, columns 1-48 |
| 10 | Time | 30 min | Incubation, luminescence evolution |
| 11 | Detection | Luminescence | ViewLux uHTS Microplate Imager (PerkinElmer) |
| Step | Notes | | |
| 1, 2, 5 | White Medium Binding 1536-well plates (white, flat bottom, medium binding Cat# AWK010000A, Aurora Microplates, Scottsdale, AZ); Reaction buffer: 50 mM Tris, pH 8.0, 3 mM MgCl_2_, 1 mM EDTA, 50 mM NaCl, 1 mM DTT, and 0.1 mg/mL BSA; refer to tables 2 and S1 for specific reagent concentrations | | |
| 3, 4 | Pintool transfer | | |
| 6, 8, 10 | Room temperature | | |
| 6 | Final reaction conditions: 20 nM PNMT, 5 µM SAM, 15 µM norepinephrine, 50 mM Tris, pH 8.0, 3 mM MgCl_2_, 1 mM EDTA, 50 mM NaCl, 1 mM DTT, and 0.1 mg/mL BSA | | |
| 8 | Conversion of SAH to ADP; MTase Glo Kit (Promega, Madison, WI) | | |
| 10 | Conversion of ADP to ATP and detection by UltraGlo luciferase; MTase Glo Kit (Promega, Madison, WI) | | |
| 11 | Settings: 20 s exposure, 1X binning, high gain, medium speed | | |

| Protocol table for NNMT HTS MTase-Glo Assay | | | |
| --- | --- | --- | --- |
| Step | Parameter | Value | Description |
| 1 | Reagent | 2 µL | NNMT (30 nM) in reaction buffer, columns 1-48 |
| 2 | Reagent | 1 µL | SAM (20 uM) in reaction buffer, columns 1-48 |
| 3 | Controls | 23 nL | DMSO in column 4; sinefungin in DMSO (0 µM – 40 µM) in 7-point 1:2 dilution series (n = 2) in column 2 |
| 4 | Library compounds | 23 nL | Columns 5-48 |
| 5 | Reagent | 1 µL | Nicotinamide (20 µM) in reaction buffer, columns 2-48 |
| 6 | Time | 20 min | Incubation |
| 7 | Reagent | 1 µL | MTase-Glo reagent (5X), columns 1-48 |
| 8 | Time | 30 min | Incubation |
| 9 | Reagent | 5 uL | MTase-Glo Detection reagent, columns 1-48 |
| 10 | Time | 30 min | Incubation, luminescence evolution |
| 11 | Detection | Luminescence | ViewLux uHTS Microplate Imager (PerkinElmer) |
| Step | Notes | | |
| 1, 2, 5 | White Medium Binding 1536-well plates (white, flat bottom, medium binding Cat# AWK010000A, Aurora Microplates, Scottsdale, AZ); Reaction buffer: 50 mM Tris, pH 8.0, 3 mM MgCl_2_, 1 mM EDTA, 50 mM NaCl, 1 mM DTT, and 0.1 mg/mL BSA; refer to tables 2 and S1 for specific reagent concentrations | | |
| 3, 4 | Pintool transfer | | |
| 6, 8, 10 | Room temperature | | |
| 6 | Final reaction conditions: 15 nM NNMT, 5 µM SAM, 5 µM nicotinamide, 50 mM Tris, pH 8.0, 3 mM MgCl_2_, 1 mM EDTA, 50 mM NaCl, 1 mM DTT, and 0.1 mg/mL BSA | | |
| 8 | Conversion of SAH to ADP; MTase Glo Kit (Promega, Madison, WI) | | |
| 10 | Conversion of ADP to ATP and detection by UltraGlo luciferase; MTase Glo Kit (Promega, Madison, WI) | | |
| 11 | Settings: 20 s exposure, 1X binning, high gain, medium speed | | |

| Protocol table for HNMT HTS MTase-Glo Assay | | | |
| --- | --- | --- | --- |
| Step | Parameter | Value | Description |
| 1 | Reagent | 2 µL | HNMT (15 nM) in reaction buffer, columns 1-48 |
| 2 | Reagent | 1 µL | SAM (20 uM) in reaction buffer, columns 1-48 |
| 3 | Controls | 23 nL | DMSO in column 4; sinefungin in DMSO (0 µM – 40 µM) in 7-point 1:2 dilution series (n = 2) in column 2 |
| 4 | Library compounds | 23 nL | Columns 5-48 |
| 5 | Reagent | 1 µL | Histamine (40 µM) in reaction buffer, columns 2-48 |
| 6 | Time | 20 min | Incubation |
| 7 | Reagent | 1 µL | MTase-Glo reagent (5X), columns 1-48 |
| 8 | Time | 30 min | Incubation |
| 9 | Reagent | 5 uL | MTase-Glo Detection reagent, columns 1-48 |
| 10 | Time | 30 min | Incubation, luminescence evolution |
| 11 | Detection | Luminescence | ViewLux uHTS Microplate Imager (PerkinElmer) |
| Step | Notes | | |
| 1, 2, 5 | White Medium Binding 1536-well plates (white, flat bottom, medium binding Cat# AWK010000A, Aurora Microplates, Scottsdale, AZ); Reaction buffer: 50 mM Tris, pH 8.0, 3 mM MgCl_2_, 1 mM EDTA, 50 mM NaCl, 1 mM DTT, and 0.1 mg/mL BSA; refer to tables 2 and S1 for specific reagent concentrations | | |
| 3, 4 | Pintool transfer | | |
| 6, 8, 10 | Room temperature | | |
| 6 | White Medium Binding 1536-well plates (white, flat bottom, medium binding Cat# AWK010000A, Aurora Microplates, Scottsdale, AZ); Reaction buffer: 50 mM Tris, pH 8.0, 3 mM MgCl_2_, 1 mM EDTA, 50 mM NaCl, 1 mM DTT, and 0.1 mg/mL BSA; refer to tables 2 and S1 for specific reagent concentrations | | |
| 8 | Conversion of SAH to ADP; MTase Glo Kit (Promega, Madison, WI) | | |
| 10 | Conversion of ADP to ATP and detection by UltraGlo luciferase; MTase Glo Kit (Promega, Madison, WI) | | |
| 11 | Settings: 20 s exposure, 1X binning, high gain, medium speed | | |

| Protocol table for GAMT HTS MTase-Glo Assay | | | |
| --- | --- | --- | --- |
| Step | Parameter | Value | Description |
| 1 | Reagent | 2 µL | GAMT (60 nM) in reaction buffer, columns 1-48 |
| 2 | Reagent | 1 µL | SAM (20 uM) in reaction buffer, columns 1-48 |
| 3 | Controls | 23 nL | DMSO in column 4; sinefungin in DMSO (0 µM – 40 µM) in 7-point 1:2 dilution series (n = 2) in column 2 |
| 4 | Library compounds | 23 nL | Columns 5-48 |
| 5 | Reagent | 1 µL | Guanidino acetate (20 µM) in reaction buffer, columns 2-48 |
| 6 | Time | 30 min | Incubation |
| 7 | Reagent | 1 µL | MTase-Glo reagent (5X), columns 1-48 |
| 8 | Time | 30 min | Incubation |
| 9 | Reagent | 5 uL | MTase-Glo Detection reagent, columns 1-48 |
| 10 | Time | 30 min | Incubation, luminescence evolution |
| 11 | Detection | Luminescence | ViewLux uHTS Microplate Imager (PerkinElmer) |
| Step | Notes | | |
| 1, 2, 5 | White Medium Binding 1536-well plates (white, flat bottom, medium binding Cat# AWK010000A, Aurora Microplates, Scottsdale, AZ); Reaction buffer: 50 mM Tris, pH 8.0, 3 mM MgCl_2_, 1 mM EDTA, 50 mM NaCl, 1 mM DTT, and 0.1 mg/mL BSA; refer to tables 2 and S1 for specific reagent concentrations | | |
| 3, 4 | Pintool transfer | | |
| 6, 8, 10 | Room temperature | | |
| 6 | Final reaction conditions: 15 nM GAMT, 5 µM SAM, 5 µM guanidino acetate, 50 mM Tris, pH 8.0, 3 mM MgCl_2_, 1 mM EDTA, 50 mM NaCl, 1 mM DTT, and 0.1 mg/mL BSA | | |
| 8 | Conversion of SAH to ADP; MTase Glo Kit (Promega, Madison, WI) | | |
| 10 | Conversion of ADP to ATP and detection by UltraGlo luciferase; MTase Glo Kit (Promega, Madison, WI) | | |
| 11 | Settings: 20 s exposure, 1X binning, high gain, medium speed | | |

| Protocol table for GNMT HTS MTase-Glo Assay | | | |
| --- | --- | --- | --- |
| Step | Parameter | Value | Description |
| 1 | Reagent | 2 µL | GNMT (30 nM) in reaction buffer, columns 1-48 |
| 2 | Reagent | 1 µL | SAM (40 uM) in reaction buffer, columns 1-48 |
| 3 | Controls | 23 nL | DMSO in column 4; sinefungin in DMSO (0 µM – 40 µM) in 7-point 1:2 dilution series (n = 2) in column 2 |
| 4 | Library compounds | 23 nL | Columns 5-48 |
| 5 | Reagent | 1 µL | Glycine (2000 µM) in reaction buffer, columns 2-48 |
| 6 | Time | 30 min | Incubation |
| 7 | Reagent | 1 µL | MTase-Glo reagent (5X), columns 1-48 |
| 8 | Time | 30 min | Incubation |
| 9 | Reagent | 5 uL | MTase-Glo Detection reagent, columns 1-48 |
| 10 | Time | 30 min | Incubation, luminescence evolution |
| 11 | Detection | Luminescence | ViewLux uHTS Microplate Imager (PerkinElmer) |
| Step | Notes | | |
| 1, 2, 5 | White Medium Binding 1536-well plates (white, flat bottom, medium binding Cat# AWK010000A, Aurora Microplates, Scottsdale, AZ); Reaction buffer: 50 mM Tris, pH 8.0, 3 mM MgCl_2_, 1 mM EDTA, 50 mM NaCl, 1 mM DTT, and 0.1 mg/mL BSA; refer to tables 2 and S1 for specific reagent concentrations | | |
| 3, 4 | Pintool transfer | | |
| 6, 8, 10 | Room temperature | | |
| 6 | Final reaction conditions: 15 nM COMT, 10 µM SAM, 500 µM glycine, 50 mM Tris, pH 8.0, 3 mM MgCl_2_, 1 mM EDTA, 50 mM NaCl, 1 mM DTT, and 0.1 mg/mL BSA | | |
| 8 | Conversion of SAH to ADP; MTase Glo Kit (Promega, Madison, WI) | | |
| 10 | Conversion of ADP to ATP and detection by UltraGlo luciferase; MTase Glo Kit (Promega, Madison, WI) | | |
| 11 | Settings: 20 s exposure, 1X binning, high gain, medium speed | | |

| Protocol table for HTS MTase-Glo Counterscreen Assay | | | |
| --- | --- | --- | --- |
| Step | Parameter | Value | Description |
| 1 | Reagent | 3 µL | SAH (1.33 uM) in reaction buffer, columns 1-48 |
| 2 | Reagent | 1 µL | SAM (16 uM) in reaction buffer, columns 1-48 |
| 3 | Controls | 23 nL | DMSO in column 4; sinefungin in DMSO (0 µM – 40 µM) in 7-point 1:2 dilution series (n = 2) in column 2 |
| 4 | Library compounds | 23 nL | Columns 5-48 |
| 5 | Time | 20 min | Incubation |
| 6 | Reagent | 1 µL | MTase-Glo reagent (5X), columns 1-48 |
| 7 | Time | 30 min | Incubation |
| 8 | Reagent | 5 uL | MTase-Glo Detection reagent, columns 1-48 |
| 9 | Time | 30 min | Incubation, luminescence evolution |
| 10 | Detection | Luminescence | ViewLux uHTS Microplate Imager (PerkinElmer) |
| Step | Notes |  |  |
| 1, 2 | White Medium Binding 1536-well plates (white, flat bottom, medium binding Cat# AWK010000A, Aurora Microplates, Scottsdale, AZ); Reaction buffer: 50 mM Tris, pH 8.0, 3 mM MgCl_2_, 1 mM EDTA, 50 mM NaCl, 1 mM DTT, and 0.1 mg/mL BSA; refer to tables 2 and S1 for specific reagent concentrations | | |
| 3, 4 | Pintool transfer | | |
| 5, 7, 9 | Room temperature | | |
| 5 | Final reaction conditions: 1 µM SAH, 4 µM SAM, 50 mM Tris, pH 8.0, 3 mM MgCl_2_, 1 mM EDTA, 50 mM NaCl, 1 mM DTT, and 0.1 mg/mL BSA | | |
| 7 | Conversion of SAH to ADP; MTase Glo Kit (Promega, Madison, WI) | | |
| 9 | Conversion of ADP to ATP and detection by UltraGlo luciferase; MTase Glo Kit (Promega, Madison, WI) | | |
| 10 | Settings: 20 s exposure, 1X binning, high gain, medium speed | | |
